# The Effect of Sudanese Smokeless Tobacco (Toombak) using on Oral Microbiota

**DOI:** 10.1101/2020.04.03.023408

**Authors:** Mudathir Abdelshafea Abdelkareem Abakar, Abdalazim Abubaker Ismail Omer, Albashir Mosab Mustafa Yousif

**Affiliations:** Assistant professor, Department of Medical Microbiology and Immunology, Alzaiem Alazhari University, Khartoum, Sudan; Department of Medical Microbiology and Immunology, Alzaiem Alazhari University, Khartoum, Sudan

**Keywords:** Smokeless tobacco, Toombak, oral Microbiota

## Abstract

**Background:** *Toombak*, *saffa* and *saod* are the common local name of Sudanese smokeless tobacco which used by large population in different regions in Sudan. That chemical nature of Toomback includes very high levels of carcinogenic substances, strong alkaline materials and microorganisms. The used of Toombak implying application of the highly addictive product in the mouth many times per day for long times differed from users to other.

**Objectives:** The objective of this study was to study the effect of Toombak on the oral microbial flora (Microbiota).

**Materials and Methods:** 100 buccal swabs were collected in this study, 50 swabs from Toombak users and 50 swabs from non Toombak users as control group. All the participants were non smokers. The swabs were cultured on Chocolate agar, Blood agar and Sabouraud agar and incubated aerobically for up to 7 days. Then colonial morphology, indirect microscopy and biochemical tests were used to identify the isolated organism.

**Results:** Toomback shows inhibitory effect on the *viridanas streptococci* and the baccual swabs collected from toomback users show significant increase in number of mixed growth (P value 0.000/ likelihood ratio 43.24) and colonization of oral cavity with *Bacillus species* (P value 0.000/ likelihood ratio 57.41), *Aspergillus species* (P value 0.003/ likelihood ratio 10.462) and *Aspergillus flavus* (P value 0.036/ likelihood ratio 4.45).

**Conclusion:** Toombak using is affecting the quality and quantity of the oral normal flora because it has inhibitory effect on *viridians streptococci*, leading to rising of new colonizing species such as *Bacillus species* and *Aspergillus specie* especially aflatoxins producing *Aspergillus flavus*. These changes in the microenvironment may affect the immunity and health of the Toombak users and may lead to serious oral and systemic health problems.

## Introduction

Using of the smokeless tobacco is common worldwide practice in more than 70 countries with different socioeconomic background especially in South-East Asia Region, which includes about 89% of the world’s users ^(1)^.

Many studies were conducted and connected the smokeless tobacco with different pathological conditions mainly cancers including oral cavity, nasal cavity, esophagus, pancreas, lung, trachea and liver in human and animal models. This wide range of cancers refers to many carcinogenic substances which counted more than 30 carcinogens have been identified in different brands of smokeless tobacco ^(1, 2, 3)^.

## Oral and systemic effect of smokeless tobacco (SLT)

The South African smokeless tobacco product contains very the high PH in the ^(4)^ which associated with development of oral keratotic lesions ^(5)^. More severe lesions have been associated with higher pH values supporting the idea that a higher product pH is associated with increased toxicity ^(6)^. Smokeless tobacco also significantly associated to oral health problems included gingival recession, leukoplakia tooth loss and loss of periodontal attachment on the side of product placement is supported in previous studies ^(7, 8, 9)^. Xerostomia, white mucosal lesions and systemic effects such as cardiovascular system, digestive and genitourinary symptoms had been reprted and related to some brands of smokeless tobacco ^(10, 11, 12, 13, 14)^.

### Toombak

The Sudanese form of smokeless tobacco product (Toombak) using in form of snuff-dipping process called (Suffa) which a small ball of Toombak commonly applied over the gums and teeth and sucked slowly for 10 to 15 minutes, the process repeated about 20 time per day in average^(15, 16)^.

Toomback is prepared from leaves of *Nicotiana rustica*by in high alkaline condition of aqueous solution of sodium bicarbonate, the process involve fermentation for up to one year and other complex process resulted in formation of moist paste form Toombak which is has strong aroma, highly addictive and contains high level of nicotine and nornicotine at least 100-fold higher concentrations of the tobacco-specific N-nitrosamines (TSNA) than US and Swedish commercial snuff brands ^(17, 18)^.

### Normal flora

Normal flora or bacterial normal flora or microflora is the population of microorganisms that inhabit the skin and mucous membranes of healthy normal persons. About half of the oral microflora are *viridians streptococci* such as *Streptococcus mutans* and *Streptococcus sanguinis*. The other half include many bacterial families such as *Eikenella corrdens, Bacteroides, provetella, Fusbacterium*, *Colstridium,* and *Peptostreptococcus* and *Actinomyces species*. This normal flora considers a first defense line against microbial pathogens, beside their role maturation of the immune system ^(19, 20)^.

### Rationale

The carcinogenic effect of Toomback has been intensively studied. The aim of this work is to study the effect of Toombak on oral microbial flora. The problem of this study rose from the hypothesis that if Toomback has an effect on the balance of microflora, this may contribute to oral, dental and periodontal infections and may affect the immunity of oral mucosa.

## Method and material

This case control study was conducted in Alzaiem Alazhari University, Khartoum, Sudan during the period from March to June 2018. The study included 100 male; 50 of them were Toombak users and they sets as case group, and the rest 50 participants were never used Toombak before their participation. All of the participants were non smokers.

Oral swabs (buccal) were collected from the entire participants for isolation and Identification of isolated organism. Antimicrobial sensitivity of Toombak was tested on the isolated organisms.

The collected swabs were cultured on blood agar, chocolate blood agar and sabouraud agar and incubated aerobically at 37ºC for 24-72 hours, with addition of 5-10% carbon dioxide enrich environment for chocolate agar.

Colonial morphology, indirect gram stain and biochemical tests were used for identification of bacterial isolates.

Needle mount with lactophenol cotton blue stain were used for identification of isolated fungi based on their morphological features.

## Sensitivity test for Toombak to isolated micro organism

Under sterile condition make wells on sterile media. Then weight the Toombak about 3g /ml of normal saline. Culture the microorganism under the test on the media which contains the wells. Add the snuff solution to the wells. Incubate at 37c for 24 hours. Measure the inhibition zone.

## Ethic consideration

This study was approved to carry out by the ethical committee of Alzaiem Alazhari University. Every participant was informed by the objective of the study, purpose of the study and received formal consent for voluntary participation, and their privacy and confidentiality have been maintained and protected.

## Statistical analysis

Frequencies, independent T-Test, Chi square test and likelihood ratio were computerizing calculated by statistical package for social science (SPSS^®^) program 21.

## Results

All of the oral swabs obtained from the 100 participants of the study, showed growth, 53(53%) of the growth was yelled one organism. The most isolated bacteria is *viridanas streptococci* which grow on the 96(96%) of the specimens. Whereas *Aspergillus species* is the only isolated fungi in this study and grow from 33(33%) specimen (Table1).

**Table (1).**
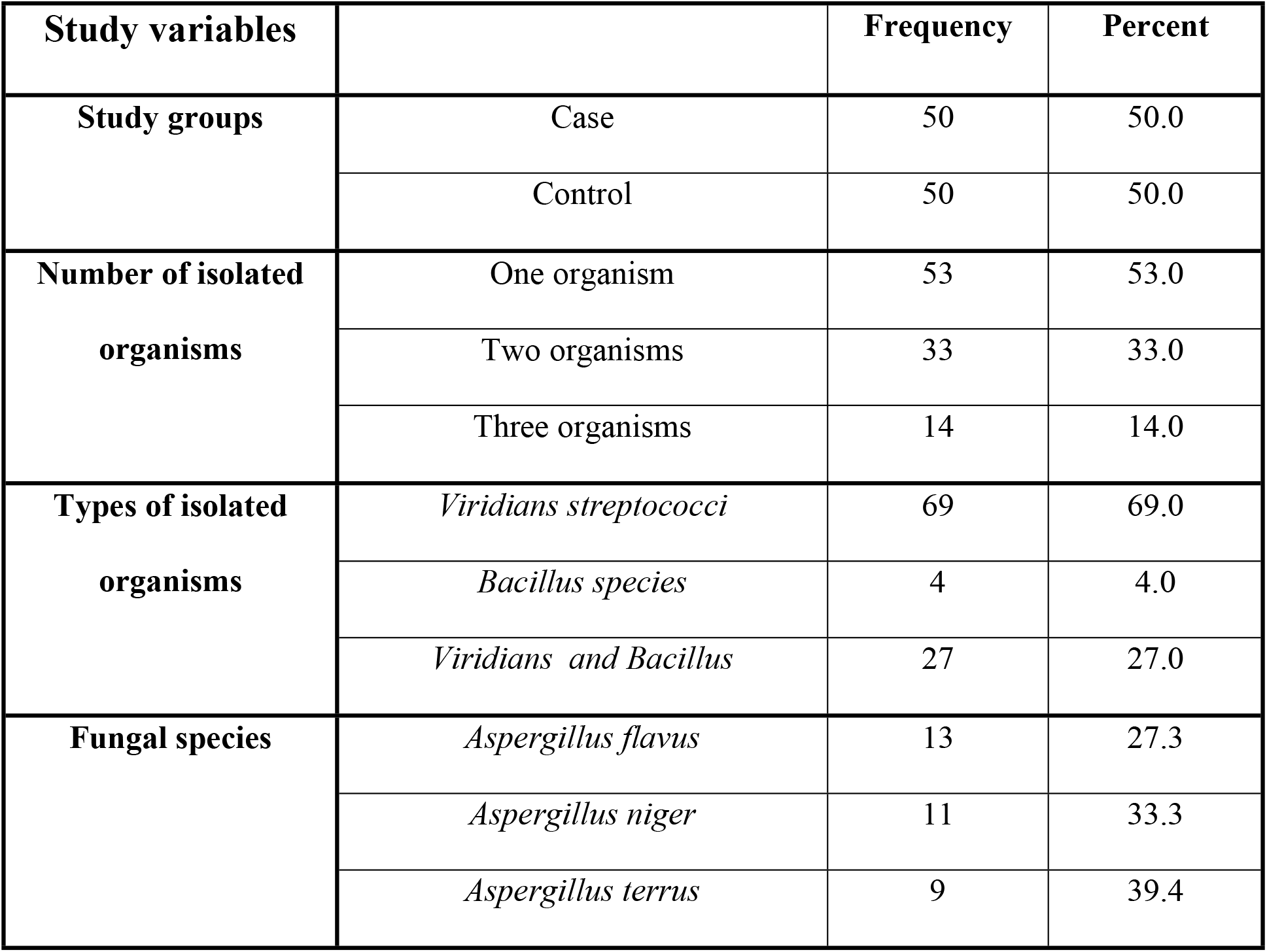
shows Study variables.

| **Study variables** |  | **Frequency** | **Percent** |
| --- | --- | --- | --- |
| **Study groups** | Case | 50 | 50.0 |
|  | Control | 50 | 50.0 |
| **Number of isolated organisms** | One organism | 53 | 53.0 |
|  | Two organisms | 33 | 33.0 |
|  | Three organisms | 14 | 14.0 |
| **Types of isolated organisms** | *Viridians streptococci* | 69 | 69.0 |
|  | *Bacillus species* | 4 | 4.0 |
|  | *Viridians and Bacillus* | 27 | 27.0 |
| **Fungal species** | *Aspergillus flavus* | 13 | 27.3 |
|  | *Aspergillus niger* | 11 | 33.3 |
|  | *Aspergillus terrus* | 9 | 39.4 |

This study revealed that there were no inhibition zone of Toombak on *Bacillus species* and *Aspergillus species*. The inhibitory effect was noted *viridanas streptococci* but there were insignificant mean differences between the mean of inhibitory zone of Toombak among *viridanas streptococci* isolated from the Toombak users and control group (Table 2).

**Table (2).**
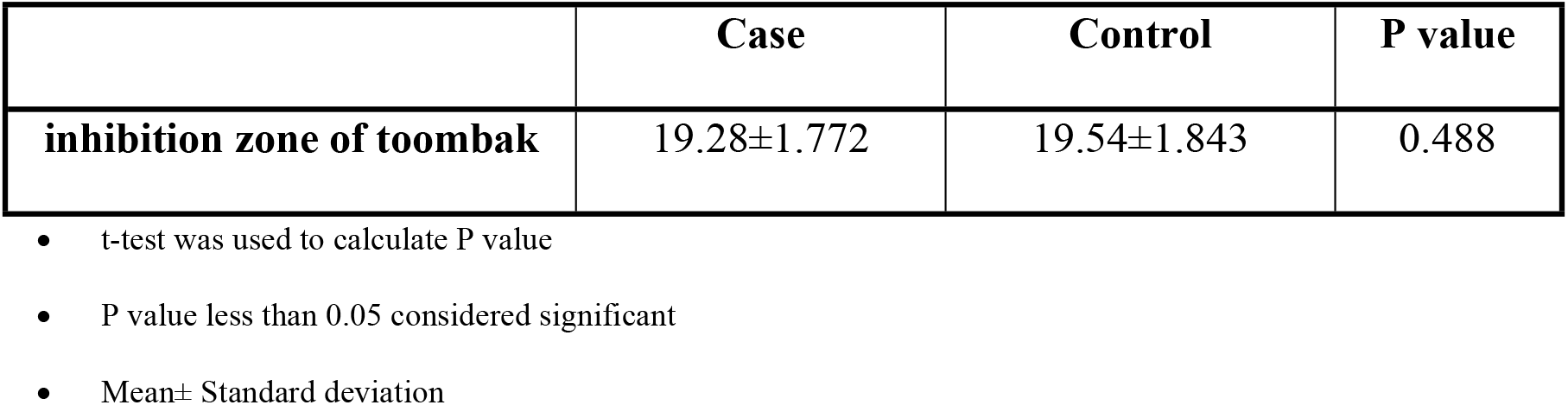
shows statistics and mean differences of inhibition zone of snuff against isolated bacteria among case and control groups.

|  | **Case** | **Control** | **P value** |
| --- | --- | --- | --- |
| **inhibition zone of toombak** | 19.28±1.772 | 19.54±1.843 | 0.488 |

- t-test was used to calculate P value - P value less than 0.05 considered significant - Mean± Standard deviation

Table 3 illustrated significant increase in number of mixed growth among Toombak users (P value 0.000/ likelihood ratio 43.24). And Toomback users have less significant frequencies of pure growth of *viridanas streptococci* (P value 0.000/ likelihood ratio 57.41). There were a significant association between using of Toombak and colonization of oral cavity with *Bacillus species* (P value 0.000/ likelihood ratio 57.41) and *Aspergillus species* (P value 0.003/ likelihood ratio 10.462). 10 out of the 13 strains of *Aspergillus flavus* (76.9%) was isolated from Toombak users, this give a significantly association of oral colonization of *Aspergillus flavus* with Toombak using (P value 0.036/ likelihood ratio 4.45).

**Table (3).** shows association of study variables with study groups.

|  |  | Study groups |  | P value | L R |
| --- | --- | --- | --- | --- | --- |
|  |  | Case | Control |  |  |
| **Number of the isolated organisms** | One organism | 12(12.0%) | 41(41.0%) | **0.000** | **43.26** |
|  | Two organisms | 24(24.0%) | 9(9.0%) |  |  |
|  | Three organisms | 14(14.0%) | 0(0.0%) |  |  |
| **Isolated organisms** | *Viridians streptococci* | 19 (19.0%) | 50(50.0%) | **0.000** | **57.41** |
|  | *Bacillus species* | 4(4.0%) | 0(0.0%) |  |  |
|  | *Viridians streptococci and Bacillus* | 27(27.0%) | 0(0.0%) |  |  |
| **Bacillus species** | **Yes** | 31(31.0%) | 0(0.0%) | **0.000** | **57.41** |
|  | **No** | 19(19.0%) | 50(50.0%) |  |  |
| **Fungi** | **Yes** | 24(24.0%) | 9(9.0%) | **0.003** | **10.462** |
|  | **No** | 26(26.0%) | 41(41.0%) |  |  |
| **Fungal species** | *Aspergillus flavus* | 10(30.3%) | 3(9.1%) | 0.708 | 0.627 |
|  | *Aspergillus terrus* | 7( 21.2%) | 2(6.1%) |  |  |
|  | *Aspergillus niger* | 7( 21.2%) | 4(12.1%) |  |  |
| *Aspergillus flavus* | **Yes** | 10(10%) | 3(35) | **0.036** | **4.54** |
|  | **No** | 40(40%) | 47(47%) |  |  |

- Chi square test was used to calculate p value - P value less than 0.05 considered significant - LR= Likelihood Ratio

## Discussion

This study was studied the effect of Toombak on oral micro-flora. At least to our knowledge there is no published study about the effect of Toombak on oral micro flora.

In this study we found significant increase in number and types of isolated microorganisms among Toombak users. Many chemicals products of microbial metabolisms was detected in tobacco and used as markers which indicate bacterial and fungal growth in ^(21, 22)^. In addition to that, the high content of different types of microorganisms such as bacteria and fungi in growing tobacco which may reach 10^5^ to 10^6^ organism per gram of leaf materials with increase in microorganism concentration by 10 to 20 times during beginning of the process ^(23)^.

Our study revealed the were no inhibitory effect of Toombak on *Bacillus species* and *Aspergillus species* with more inhibitory effects on *viridians streptococci*, beside a significant decrease in number of pure *viridians streptococci* isolates among Toombak users. In the other hand, we found that there was significant association between using of Toombak and colonization of oral mucosa with *Bacillus species* and *Aspergillus species*. Several studies were enumerating the bacterial populations of tobacco with predominant presence of *Bacillus species* and *Aspergillus species* ^(24,25, 26, 27, 28, 29, 30, 31,)^. This may be due to presence of spores which make the *Bacillus species* more resistant to harsh environment following fermentation of tobacco ^(24, 25, 27)^.

Regarding the inhibitory effect against *viridians streptococci*, this finding is similar to study conducted by *Qandil R* ^(32)^ show that *viridians streptococci* is more susceptible to the effects of nicotine in contrast to *staphylococcus aureus*, *spirocheate and B.burgdorferi* were only slightly inhibited or were completely unaffected following exposure to nicotine. Another invitro study conducted by *Lindemeyer RG* ^(33)^ shows contrast to our study which show stimulation of the growth of *Streptococcus mutans* and *Streptococcus sanguis* in the presence of smokeless tobacco extracts this difference may be due to difference in sugar and nicotine content of the smokeless tobacco product.

The present study shows a significant association between oral colonization with *Aspergillus flavus* and using of Toombak. This agreed with the work of *Varma SK* ^(34)^ and his accompaniments who reported presence of aflatoxins producing *Aspergillus* in the tobacco.

## Conclusion

Toombak using is affecting the quality and quantity of the oral normal flora because it has inhibitory effect on *viridians streptococci*, leading to rising of new colonizing species such as *Bacillus species* and *Aspergillus specie* especially aflatoxins producing *Aspergillus flavus*. These changes in the microenvironment may affect the immunity and health of the Toombak users and may lead to serious oral and systemic health problems.

